# Multiplex generation, tracking, and functional screening of substitution mutants using a CRISPR/retron system

**DOI:** 10.1101/2020.01.02.892679

**Authors:** Hyeonseob Lim, Soyeong Jun, Minjeong Park, Junghak Lim, Jaehwan Jeong, Ji Hyun Lee, Duhee Bang

## Abstract

We developed a clustered regularly interspaced short palindromic repeats (CRISPR)/retron system for multiplexed generation of substitution mutations by co-utilization of a retron system that continuously expresses donor DNA and a CRISPR/Cas9 cassette that induces cleavage at target genomic loci. Our system efficiently introduces substitution mutation in the *Escherichia coli* genome in a high-throughput manner. These substitution mutations can be tracked by analysis of retron plasmid sequences without laborious amplification of individual edited loci. We demonstrated that our CRISPR/retron system can introduce thousands of mutations in a single experiment and be used for screening phenotypes related to chemical responses or fitness changes. We expect that our system could facilitate genome-scale substitution screenings.

## INTRODUCTION

Several genome-wide pooled screening methods have been recently developed^*1–3*^ using the CRISPR/CRISPR-associated protein-9 (Cas9) system^*4–6*^ for targeted gene knock-out, repression^*1*^, or activation^*2*^. These screening technologies utilize Cas9 and single-guide RNA (sgRNA) libraries constructed as lentiviral delivery systems that are integrated into the genome. Genome-integrated sgRNA sequences are then analyzed using next-generation sequencing (NGS) technologies to track the mutations introduced across genomic loci. The experimental procedures used in these approaches are convenient, enabling the analysis of selected mutants using algorithms^*7, 8*^. As such, these technologies have been used in a number of recent studies^*1, 2, 9*^.

The development of pooled screening technologies for substitution mutations has not advanced as rapidly as knock-out and transcriptional regulation methods. The development of pooled screening technologies is complicated by the requirement for an additional component, an independent donor DNA fragment that serves as a template for homology-directed repair and must be delivered with the corresponding sgRNA. To overcome this challenge, a recently developed Cas9 retron precise parallel editing via homology (CRISPEY)^*10*^ method uses the retron system^*11, 12*^ for donor DNA expression instead of separate transfection of donor DNA fragments. The retron system for *Escherichia coli*^*13, 14*^, comprising msr, msd, and a reverse transcriptase (RT), was modified to express single-stranded donor DNAs in yeast, and the modified retron was flanked by sgRNA. CRISPEY thus enables the preparation of both sgRNA and donor DNA using a single vector. The vector was also designed to insert a donor DNA–sgRNA pair for each target, which were prepared from commercially available oligo pools. The CRISPEY system enables high-throughput editing in which the efficiency is enhanced by the killing of unmodified cells. In addition, the abundance of each mutant can be monitored by sequencing of the vector amplified using common primers. However, although *E. coli* is the most widely used model organism^*15, 16*^ in research related to metabolic engineering, molecular biology, and therapeutics, the CRISPEY system has not been extended to editing of the *E. coli* genome.

For editing the *E. coli* genome, lambda red recombination–based methods^*17*^ such as multiplex automated genome engineering (MAGE)^*18*^ have been used to generate genomic substitutions using only a library of oligonucleotides. However, tracking genomic mutations introduced using MAGE becomes more difficult as the number of engineered target loci increases, because target loci must be analyzed individually via laborious amplification^*19*^ and subsequent sequencing.

In contrast to the above-mentioned approaches, the trackable multiplex recombineering method^*20*^ enables tracking of variants via microarray-based capture. However, this method is not appropriate for introducing substitution mutations in coding regions because a tag sequence must be inserted into the targeted region. As such, methods that enable trackable, scarless introduction of substitution mutations in *E. coli* are needed. To fill this demand, we developed a method combining the retron and CRISPR systems and demonstrated genome engineering and screening in *E. coli*^*6, 21*^ (**Figure 1a**). Our system enables highly efficient genome editing, and the abundance of each mutant is correlated with the retron portion of the plasmid; thus, mutations across the genome can be tracked using single-amplification reactions with common primers (**Figure 1b**). Finally, we demonstrated that our system is suitable for genome-scale experiments involving *E. coli*.

**Figure 1.**
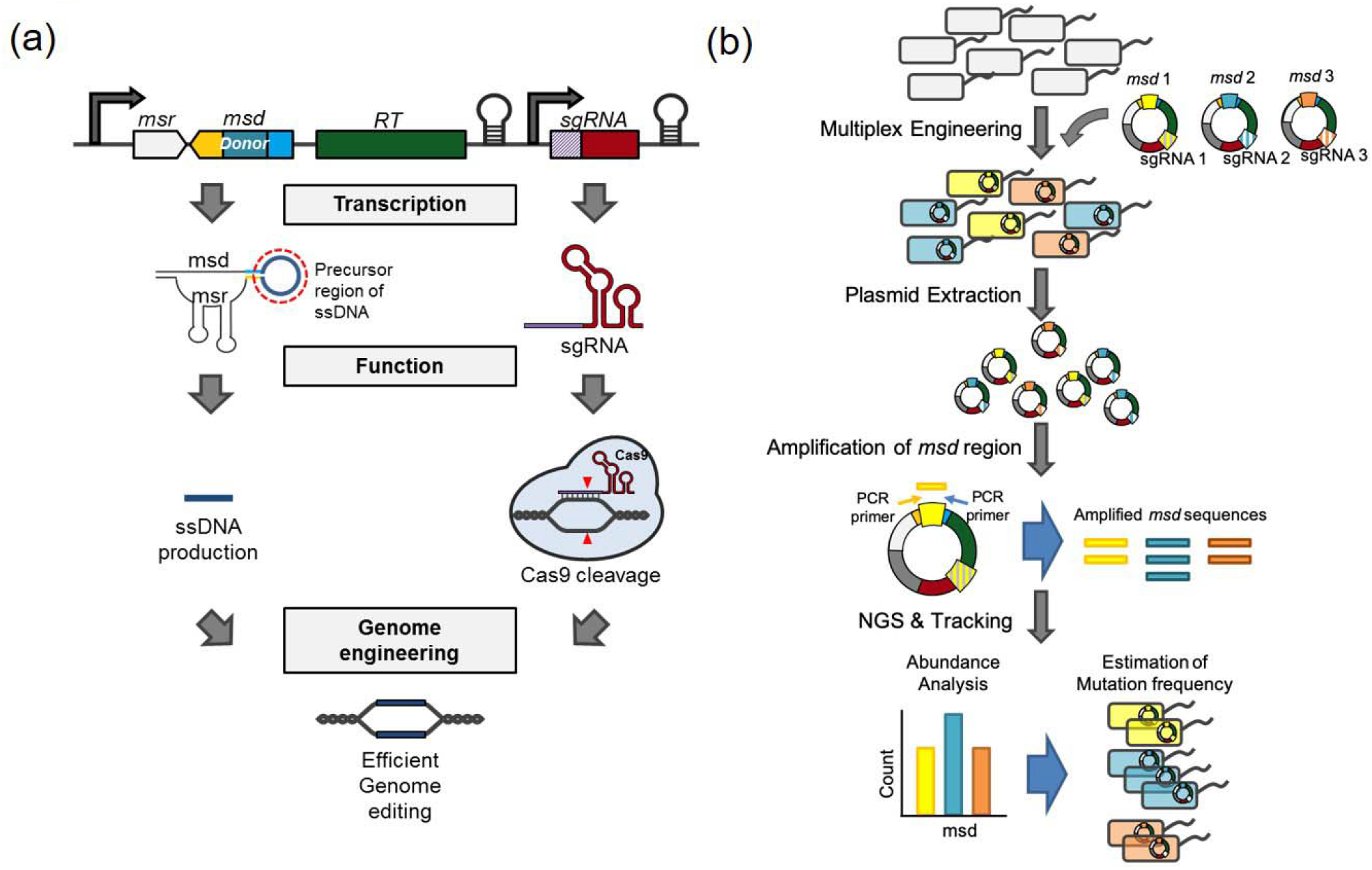
Overview of the CRISPR/retron strategy. (a) General scheme of genome editing using a CRISPR/retron cassette comprised of retron and sgRNA components. Donor ssDNA is produced from an RNA transcript containing msr and msd via reverse transcription. Cleavage of the unedited genome is induced by sgRNA and orthogonally expressed Cas9 protein. (b) Schematic workflow of *msd*-mediated variant tracking.

## RESULTS and DISCUSSION

First, we designed a programmable plasmid (pRC_blank_01) containing a sgRNA-expressing region and retron-encoding region, with the latter composed of the *msd*, *msr*, and *RT* (**Supplementary Figure S1a**, **Tables S1 and S2**). We then induced a single amino acid substitution to revert a premature stop codon (I239*) in the galactokinase enzyme (encoded by *galK*) in ENC (*E. coli* EcHB3 bearing plasmids encoding T7 polymerase and Cas9) (**Figure 2a**). The 75-bp msd and sgRNA regions were cloned into pRC_blank_01 to generate pRC_*galK*_on. We then transformed pRC_*galK*_on into ENC cells via electroporation (**Figure 2b**, **Supplementary Table S3**). Transformants were cultured in medium supplemented with isopropyl β-D-1-thiogalactopyranoside (IPTG) and inoculated into fresh medium every 12 h for 15 dilution cycles. The recombination efficiency was determined at six time points (cycles 0, 3, 6, 9, 12, and 15) via NGS; we obtained a recombination efficiency of 5.7% with 15 cycles of dilution (n=3). Moreover, the engineering efficiency increased as the dilution cycles progressed, whereas deactivated retrons (dead RT) did not promote such efficiency (n=2) (**Figure 2c**).

**Figure 2.**
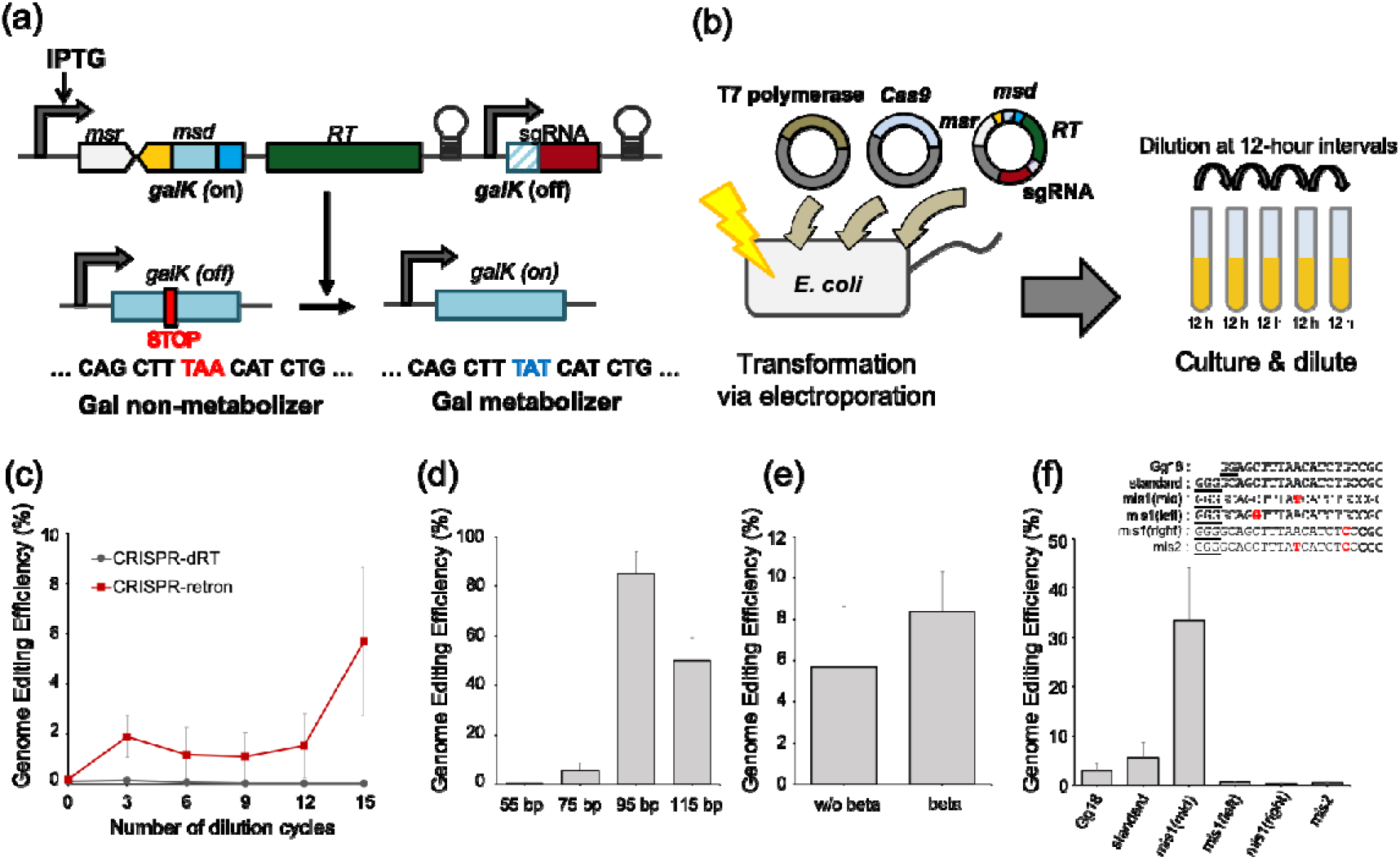
Single-plex engineering procedure and optimization. (a) Schematic workflow of *galK* reversion experiments. (b) Schematic diagram of *galK* engineering mediated by the CRISPR/retron system. (c) Relationship between engineering efficiency and dilution cycle progression. Red solid line with squares denotes the engineering efficiency of the CRISPR/retron system (n=3), and black dotted line with X marks denotes the engineering efficiency of deactivated retrons with dead RT (n=2). (d-f) Conditional dependence of CRISPR/retron on *msd* length (d), presence or absence of Beta recombinase (e), and sgRNA activity (f). Spacer sequences are shown, and guanines essential for transcription are underlined. Each value was determined from three independent experiments. Error bars indicate standard error of the mean (SEM).

Next, we tested various derivatives of pRC_*galK*_on (**Supplementary Table S2**) to optimize the system parameters. First, we varied the conditions that affect the efficiency of homologous recombination. The effects of changing the length of msd (from 55 to 115 bp)^*18, 22*^ and the presence/absence of Beta recombinase^*13, 14*^ (which stabilizes ssDNA) were tested; optimal efficiency was obtained with an msd length of 95 bp and co-expression of Beta recombinase (**Figure 2d, e**). We then examined the effect of varying conditions related to selection pressure using the CRISPR/Cas9 system. We changed the activity of the sgRNA by altering the 5’ sequence^*23*^ and the number of mismatches^*5*^ that are known to affect CRISPR/Cas9 efficiency. The sgRNA in our system is designed to have extra guanines (GGG + 20 nt) in the 5’ sequence due to the characteristics of the T7 promoter. We tested “Gg18”^*23*^, which uses a shortened T7 transcription start site “GG” as first two bases in the 20-nt spacer sequence (GG + 18 nt) to remove the 5’ extra bases. We also tested the effect of mismatches in the sgRNA sequence to modulate the efficiency of CRISPR/Cas9^*5*^. One mismatch in the middle of GGG + 20 nt formed a sequence with moderate sgRNA activity and enhanced editing efficiency (**Figure 2f**). Standard and Gg18 sgRNAs appeared to kill the cells before editing due to high CRISPR/Cas9 efficiency, and sgRNAs with mismatches introduced into the left or right side did not effectively kill unedited cells. Thus, editing efficiency was enhanced with a 95-bp msd, co-expression of Beta recombinase, and use of a sgRNA with a mismatch in the middle. To exploit these optimized conditions, we constructed an improved programmable plasmid by introducing the Beta recombinase gene into pRC_blank_01 (pRC_blank_02) (**Supplementary Figure S1b**).

The scalability of the CRISPR/retron system was assessed using degenerate bases. We also investigated whether the plasmid *msd* frequency is representative of the corresponding genomic mutant allele frequency. A *msd* oligomer containing six consecutive degenerate bases (N6) was synthesized to produce mismatch changes in the *galK* gene, and the oligomer was cloned into the plasmid with the *galK*-targeting sgRNA (**Figure 3a**). We observed a recombination efficiency of 6.57% with 15 cycles of dilution (n=2) (**Supplementary Figure S3a**). The genomic engineering efficiency also increased with additional dilution cycles. On average, the engineered cell populations contained 1,651 genomic variant sequences after 15 cycles of dilution (n=2) of the 4,095 *msd*s identified from the plasmid pool before transformation (**Supplementary Figure S2**). Importantly, a strong positive correlation was observed at 15 cycles of dilution between *msd* (from a plasmid) and the genomic variant ratio based upon a comparison of normalized depth (ND) values (**Figure 3b**) (Pearson’s ρ=0.87, 0.84 for each replicate). In addition, the strength of the correlation increased with additional dilution cycles, as did clonal bias (**Supplementary Figure S3b and c**). We presume that the clonal bias was associated with differences in survival rate between mutants. Because the editing efficiency is dependent on the number of mismatches^*18*^, unedited cells containing low-efficiency donor DNA are more likely to be eliminated via selection. For further validation, we analyzed the genomes and *msd* regions of individual colonies picked from each pool using Sanger sequencing, and this analysis exhibited similar trends (**Supplementary Table S4**). Collectively, these results indicate that the variant ratio can be determined by tracking plasmid-borne *msd* sequences.

**Figure 3.**
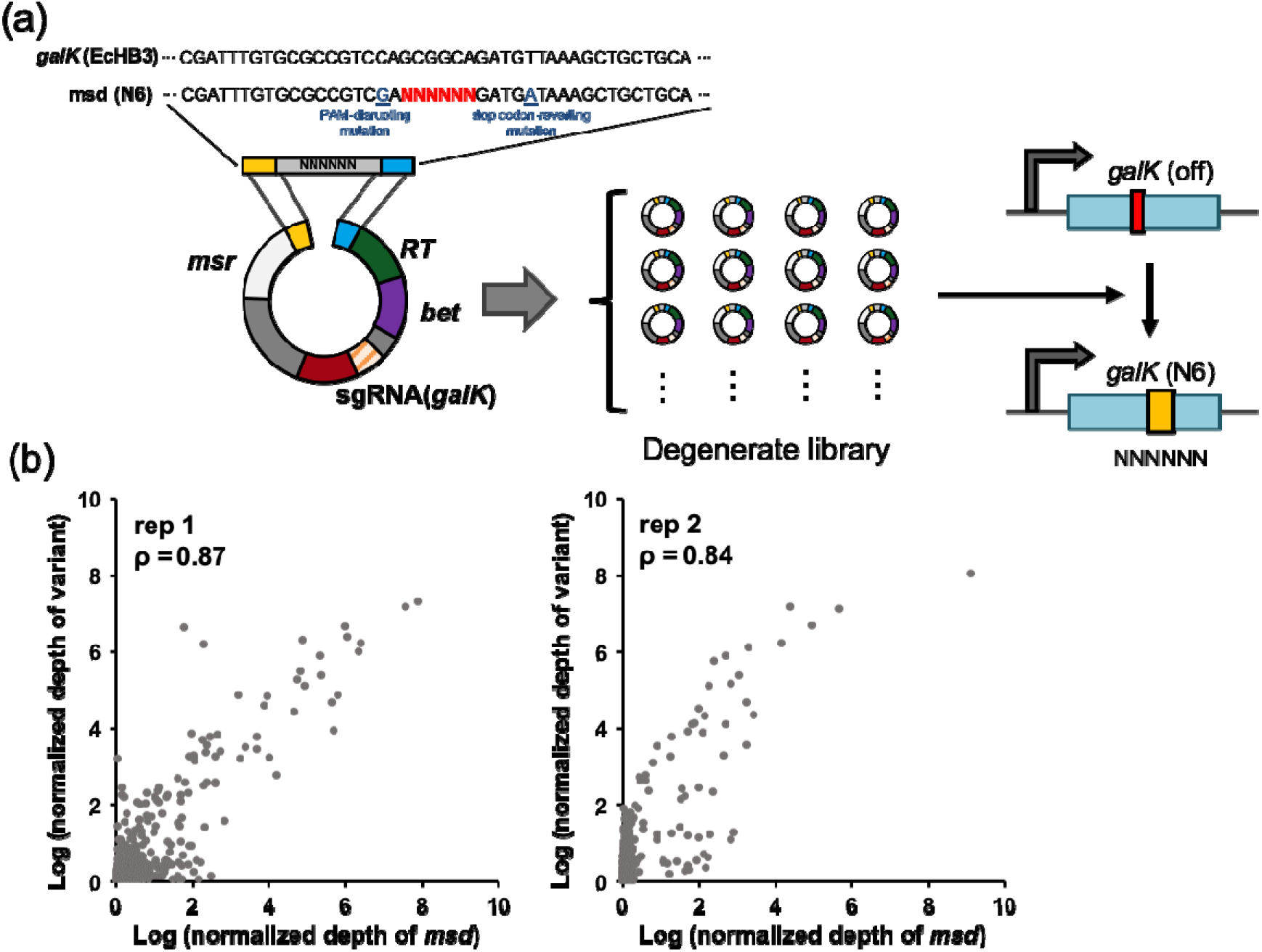
Degenerated library experiment. (a) Schematic illustration of degenerate library construction. (b) Scatter plots showing the correlation between each variant and *msd* depth. Each axis was scaled as the logarithm of normalized depth value, as described in the Methods.

Next, we conducted a model study to evaluate the CRISPR/retron system for multiplex engineering. We introduced substitution mutations in six non-essential genes (*apt*, *cadA*, *kefB*, *malK*, *rpoB*, and *tolC* [**Supplementary Table S5**]) by co-transforming ENC cells with individually constructed plasmids (**Supplementary Figure S4a**). The *msd* ratio for the plasmid of each variant was highly correlated with the genomic variant ratio at 15 cycles of dilution (**Supplementary Figure S4b**) (ρ=0.96, n=3), and the recombination rate was high (46.8%) (**Supplementary Figure S4c**). These results demonstrate that the CRISPR/retron system is suitable for use in generating and tracking substitution mutations at multiple loci.

We also investigated whether the CRISPR/retron system could be used to screen for chemical-resistant mutants (**Figure 4a**), targeting seven genes (*apt*, *cadA*, *cat*, *galK*, *kefB*, *malK*, and *rpoB* [**Supplementary Table S5**]); the *cat* product detoxifies the antibiotic chloramphenicol (cam). We designed the cat-targeted msd to revert the stop codon in the EcHB3 strain (see Methods, **Supplementary Tables S3** and **S6**); therefore, only cells carrying the cat-reverting mutation could survive. Indeed, the results of NGS analysis of genomic DNA and the *msd* region showed that only cat-reverting mutants were enriched, based upon comparisons of the ND of cam-treated versus untreated control cells among the seven mutants (**Figure 4b-d**, **Supplementary Tables S7-9**). The enrichment was also confirmed by Sanger sequencing (**Supplementary Table S10**). These results demonstrated that our CRISPR/retron system can be used to identify chemical-resistant mutants.

**Figure 4.**
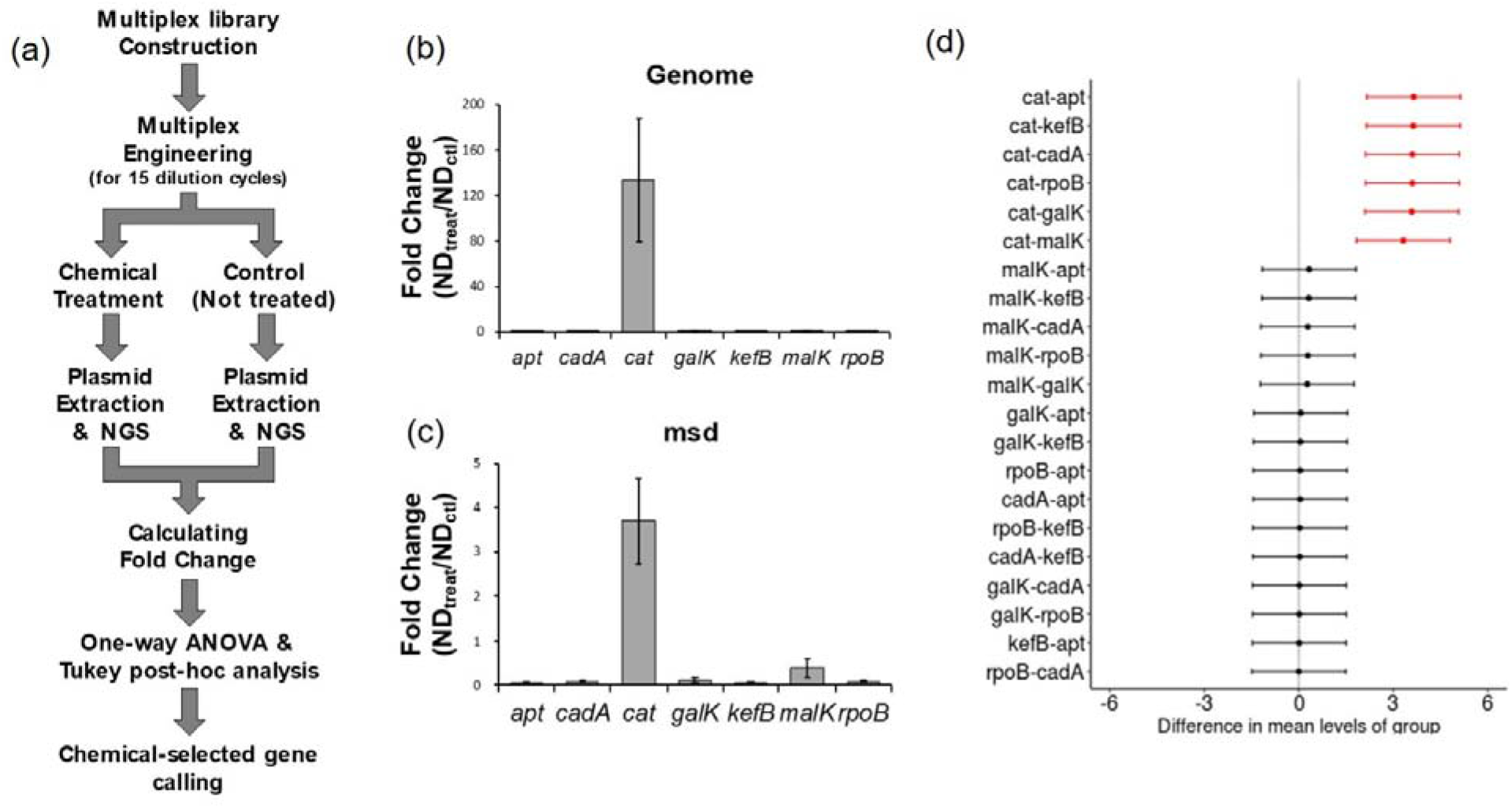
Chemical selection experiment. (a) Schematic flow diagram for detecting a chemical-selected gene. Fold-change of the genome (b) and plasmid (c) as calculated in the chloramphenicol-treatment experiment. Error bars indicate standard error of the mean (SEM). (d) Differences in mean fold-change with corresponding 95% confidence intervals obtained by Tukey post-hoc analysis.

Finally, we extended the size of the library to a genome-wide scale to facilitate screening thousands of mutations across the genome. The library was designed to enable gene knock-out by introducing a stop codon in 3,222 genes in the *E. coli* genome. *msd* and sgRNA sequences were synthesized via high-throughput oligo synthesis and inserted into vectors for transformation of *E. coli* cells (Methods, **Supplementary Figure S5**, **Table S11**) using 20 biological replicates. After 15 cycles of culture, *ygbT* was consistently enriched in a majority of the replicates. An ordinary least square model revealed that only the knock-out of *ygbT* increased in a linear manner depending on the culture period (**Supplementary Figure S6a**, **Supplementary Table S12**). The change in fitness resulting from the knock-out of *ygbT* was confirmed by constructing and transforming cells with a single plasmid targeting *ygbT* (**Supplementary Figure S6b**).

In summary, the CRISPR/retron system we developed efficiently generates programmed substitutions and enables the identification of mutants exhibiting phenotypes related to targeted chemical responses or enhanced proliferation. The feasibility of the method was demonstrated in an experiment using a genome-scale library, which identified *ygbT* as a gene involved in regulating the growth of *E. coli*, although the underlying mechanism remains to be elucidated. Our CRISPR/retron method relies on the abundance of *msd* sequences rather than the mutant allele frequency obtained by amplification of individual loci; therefore, our method is suitable for analyzing thousands of mutants. It is generally difficult to extend the size of the library using the other method^*18, 22*^, which uses individually amplified genomic sequences, because it is difficult to identify a mutation.

The CRISPR/retron system described here provides enhanced editing efficiency of the bacterial genome compared with previous methods utilizing the retron system^*13*^; however, a non-negligible proportion of unedited cells was still observed. Unedited cells can be problematic because their presence can interfere with interpretation when the screening process is not based on chemical selection. Enhancing the recombineering machinery via knock-out of exonucleases^*14*^, for example, could help reduce the proportion of unedited cells.

Although the use of our system is currently limited to bacterial engineering, we expect that it could be applied to a variety of other species by adapting their expression system, similar to the case of CRISPEY system. We anticipate that use of a lentivirus vector would yield similar results in mammalian cells, but this would require development of a chemical- or siRNA-mediated strategy to inhibit non-homologous end joining repair. We believe that integration of the above-mentioned improvements in our CRISPR/retron system would facilitate its use in the explorations of cancer-associated substitution variants, Mendelian-inherited disorders, and other phenotypes of interest in mammalian cells.

## METHODS

### Bacterial strains, antibiotics, induction, and electroporation

EcNR2^*18*^, EcHB3 (**Supplementary Table S6**), C2566 (New England BioLabs, USA), and their derivatives were grown at 30°C (37°C for C2566) in Luria-Bertani (LB) broth (or agar) supplemented with appropriate antibiotics: ampicillin, 50 μg/ml; or spectinomycin, 100 μg/ml. IPTG was added to a final concentration of 1 mM. Electroporation was performed as follows: cells were grown until reaching an OD_600_ of 0.7. Next, cells equivalent to 1 ml of culture medium were washed twice with 1 ml of ddH2O and resuspended in 50 μl of the appropriate product. The cells were then pulsed with 1.8 kV in a 1-mm-gap cuvette (Bio-Rad, USA).

### CRISPR/retron plasmid construction for targeting single genes

Oligonucleotides for constructing the *msd* and spacer inserts were designed and synthesized with the desired mutation sequences and one synonymous mutation in the protospacer adjacent motif (PAM) sequence in order to prevent killing of the engineered cells (IDT, USA) (**Supplementary Table S6**). The inserts were amplified as follows: 0.1 pmol of each template was diluted with ddH2O up to 8 μl and mixed with 1 μl of forward primer (10 μM), 1 μl of reverse primer (10 μM), and 10 μl of 2× KAPA HiFi HotStart Ready Mix (KAPA Biosystems, USA). The following conditions were used: 95°C for 3 min, 25 cycles of 95°C for 30 s, 60°C for 30 s, and 72°C for 30 s, with a final extension at 72°C for 1 min. Each *msd* and spacer insert was cloned into *Eco*RI- and *Afl*II-digested pRC_blank_01 or pRC_blank_02 vector by Gibson assembly. Information regarding the constructed plasmids and oligomer sequences is provided in **Supplementary Tables S1**, **S2**, and **S6**.

### Preparation of the degenerate library

Oligonucleotides for the N6 experiment were synthesized at IDT (**Supplementary Table S6**). The *msd* insert was constructed using PCR amplification: 1 μl of 0.1 μM *galK*_N6_oligo, 1 μl of forward primer (10 μM), 1 μl of reverse primer (10 μM), 7 μl of ddH2O, and 2× KAPA HiFi HotStart Ready Mix were combined and mixed, and reaction was performed as follows: 95°C for 3 min, 25 cycles of 95°C for 30 s, 60°C for 30 s, and 72°C for 30 s, with final extension at 72°C for 1 min. Products were purified using Ampure XP beads (Beckman Coulter, USA), secondarily amplified over eight cycles using 30-mer Gibson homology-flanked primers, and then purified again. A total of 13 ng of the product was cloned into 320 ng of the *Eco*RI-digested pRC_*galK*_sg_only vector (**Supplementary Table S6**). The plasmid was purified using Ampure XP beads and subsequently transformed into C2566 by electroporation. Cells were recovered after culture in 3 ml of LB broth overnight at 37°C. The recovered cells were added to 7 ml of LB medium and incubated overnight at 37°C. Plasmid DNA was then isolated from the transformants using an Exprep plasmid SV kit (GeneAll, Korea).

### Preparation of the genome-scale library

Spacers and donor DNAs were designed based on the MG1655 reference genome (ASM584v2). All possible spacers with PAM (NGG/NAG) sequences within the gene were listed. Then, we chose spacers that contained targetable codons that could be changed to stop codons by replacing one or two bases (i.e., NAG, NGA, NAA, TNG, TGN). We then picked one spacer per gene for which the position was between residues corresponding to 30-70% of an open reading frame. The donor DNA was designed to replace the target codon with a stop codon and contain an altered PAM sequence to prevent recognition of the edited genome.

The sequence was designed as follows in order to reduce the oligonucleotide length: GAGTTACTGTCTGTTTTCCTG-[Donor DNA]-CAGGAAACCCGTTTTTTCT-[*Eco*RI]-TAATACGACTCACTATAGGG-[Spacer]-GTTTTAGAGCTAGAAATAGCAAG. We designed the complete sequence so that it could be constructed by secondary insertion of RT-beta fragment in the *Eco*RI site after insertion of the short fragment into pRC_blank_02. In addition, the length of the donor DNA was adjusted to 91 nt so that the total sequence would not exceed 200 nt for reasons of quality and cost.

The designed oligo pool was synthesized by Twist Bioscience (US) and amplified using chip_fwd and chip_rev primers under the following conditions: 95°C for 3 min, 20 cycles of 95°C for 30 s, 55°C for 30 s, and 72°C for 30 s, with final extension at 72°C for 1 min. The resulting product was purified and secondarily amplified using chip_2nd_fwd and chip_2nd_rev. The conditions for the second PCR were as follows: 95°C for 3 min, 10 cycles of 95°C for 30 s, 55°C for 30 s, and 72°C for 30 s, with final extension at 72°C for 1 min. The amplicons were then cloned into the pRC_blank_02 plasmid using 12 parallel Gibson assembly reactions to ensure the proper library size. The product was purified and transformed into C2566 via electroporation, and the plasmid library was extracted using a Qiagen plasmid maxi prep kit. The RT-beta sequence (**Supplementary Table S1**) was inserted into this plasmid library via *Eco*RI linearization and Gibson assembly to generate the complete knock-out library. This step was performed in 12 parallel reactions.

### Genome editing using the CRISPR/retron system

EcHB3 cells were sequentially electroporated with pN249 and pCas9_T7 to express T7 polymerase and Cas9, respectively, yielding ENC cells. The CRISPR/retron plasmid was then electroporated into the ENC cells, which were then transferred to LB broth without antibiotics for a 6-h recovery. Transformants were transferred to LB broth containing appropriate selective antibiotics, incubated at 30°C, and diluted 1:100 with 10 ml of fresh LB broth every 12 h.

### Deep sequencing for genome and *msd* region

Genomic and plasmid DNA were extracted from transformants using a DNeasy Blood & Tissue kit (Qiagen, Germany) and Exprep plasmid SV kit, respectively. Targeted loci and *msd* regions were amplified from a mixture of 200 ng of template, 10 pmol of each primer pair (**Supplementary Table S6**), and 10 μl of KAPA HiFi HotStart Ready Mix. Reactions were carried out as follows: 95°C for 3 min, 25 cycles (20 cycles for *msd*) of 95°C for 30 s, 60°C for 30 s, and 72°C for 30 s, with final extension at 72°C for 1 min. In cases of more than one condition, the samples were amplified using barcoded primers and then pooled. The amplified products were prepared for NGS using a SPARK DNA Sample Prep kit (Enzymatics, USA). Sequencing was performed using an Illumina HiSeq 4000 or NextSeq 500 sequencer (data deposited in the SRA [PRJNA598315]). Raw data were demultiplexed by flanked barcodes; *msd* and targeted sequences were analyzed by comparison with the wild-type sequences.

### Correlation coefficient calculation

To investigate the correlation between *msd* and variant depth, Pearson correlation coefficients were calculated, as follows. To normalize differences in the amount of data, we corrected for depth value by normalizing the total depth to 10^4^, and 1 was added to the ND to avoid a ND of <1. This approach, also used previously^*3*^, is represented by the following equation:

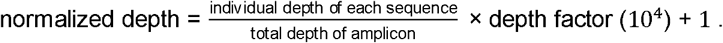

The depth factor was determined by the quantity of data. When testing the response of chloramphenicol, 10^6^ was used for the depth factor in consideration of the large amount of data. Then, the normalized *msd* depth and normalized variant depth were used to calculate the correlation coefficient.

### Chemical-resistance analysis

The significance of resistant mutant enrichment was confirmed by one-way ANOVA & Tukey post hoc analysis using the fold-change in the ND of chemical-treated samples relative to control samples.

### Investigation of ygbT knock-out on fitness

After 15 cycles of culture of ygbT-targeting plasmid transformants, the cells were cultured overnight and then diluted 1:1000 in fresh medium and incubated at 30°C. The OD_600_ was then measured every 2 h. The pRC_blank_02 plasmid was used as a control.

## AUTHOR INFORMATION

### Corresponding Authors

Address correspondence to Duhee Bang or Ji Hyun Lee.

### Author Contributions

H.L., S.J., D.B., and J.H.L. designed the study. H.L., S.J., M.P., and J.L. performed the experiments. H.L. and S.J. analyzed the data. J.J. provided the vector and advised on experimental design. H.L. and S.J. prepared the initial manuscript. D.B. and J.H.L. supervised the study and revised the final manuscript.

### Notes

D.B., S.J., and H.L. are authors of a patent application for the method described in this paper (Generation and tracking of substitution mutations in the genome using a CRISPR/retron system, KR.10-1922989). The remaining authors declare no competing financial interests.

## ACKNOWLEDGMENT

This work was supported by: (i) the Mid-career Researcher Program (NRF-2018R1A2A1A05079172), (ii) the Bio & Medical Technology Development Program (NRF-2016M3A9B6948494), (iii) the Bio & Medical Technology Development Program (NRF-2018M3A9H3024850), and (iv) Basic Science Research Program (NRF-2018R1A2B2001322) of the National Research Foundation of Korea, funded by the Ministry of Science, ICT & Planning, and (v) the Korea Health Technology R&D Project (HI18C2282) through the Korea Health Industry Development Institute (KHIDI), funded by the Ministry of Health & Welfare.

