## Supplementary Data for "Multiplex generation, tracking, and functional screening of substitution mutants using a CRISPR/retron system"

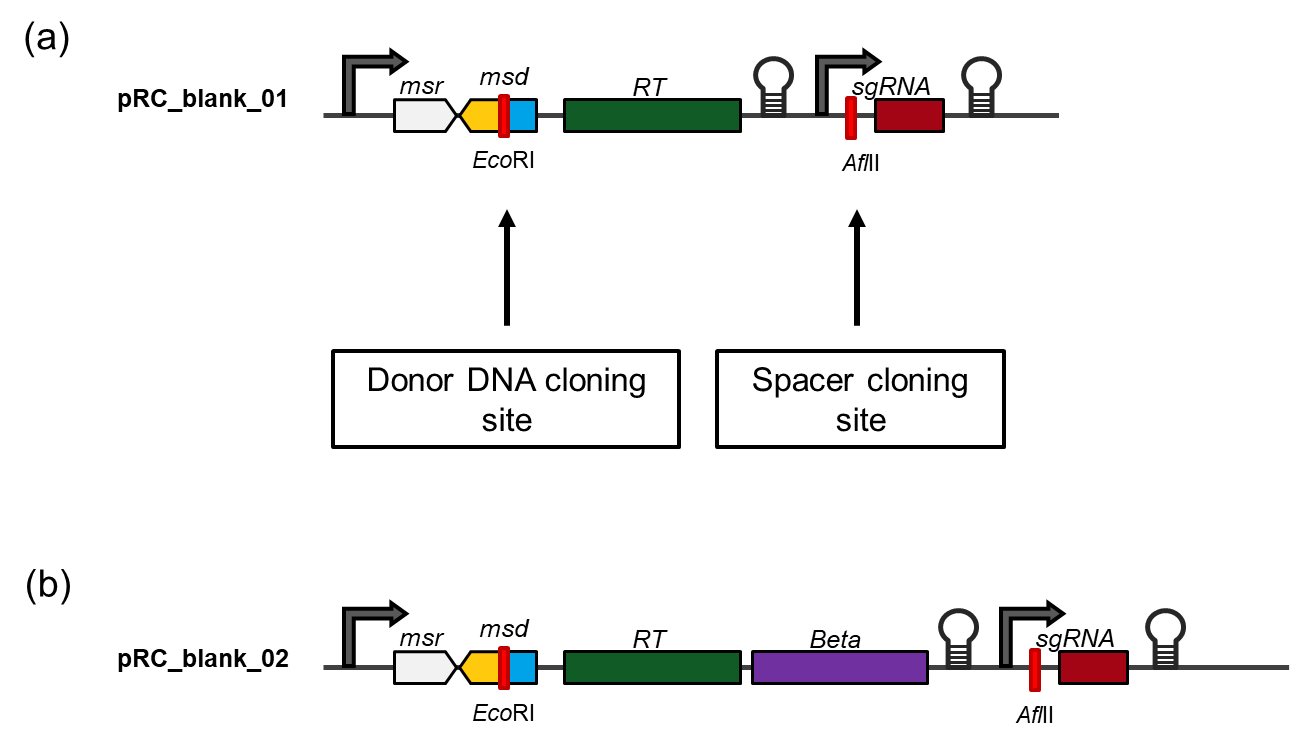

**Figure S1** | Design of the CRISPR/retron cassette used in this study. (a) Cloning site of the pRC_blank_01 plasmid. An *Eco*RI site was inserted into the *msd* sequence for cloning donor DNA, and an *Afl*II site was inserted into the sgRNA sequence for cloning spacer sequences. (b) The gene encoding Beta recombinase was inserted downstream of the reverse transcriptase gene (*RT*), yielding pRC_blank_02.

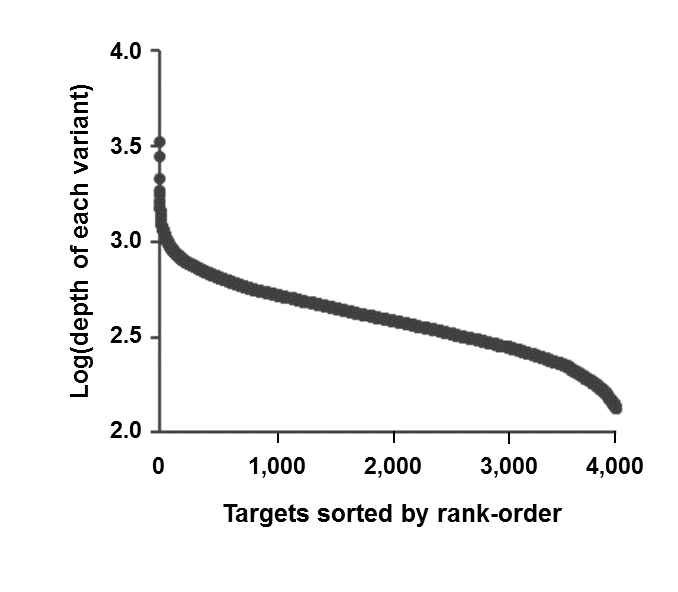

**Figure S2** | Plasmid analysis of the degenerate library. Distribution of each degenerate base sequence (N6) at plasmid construction.

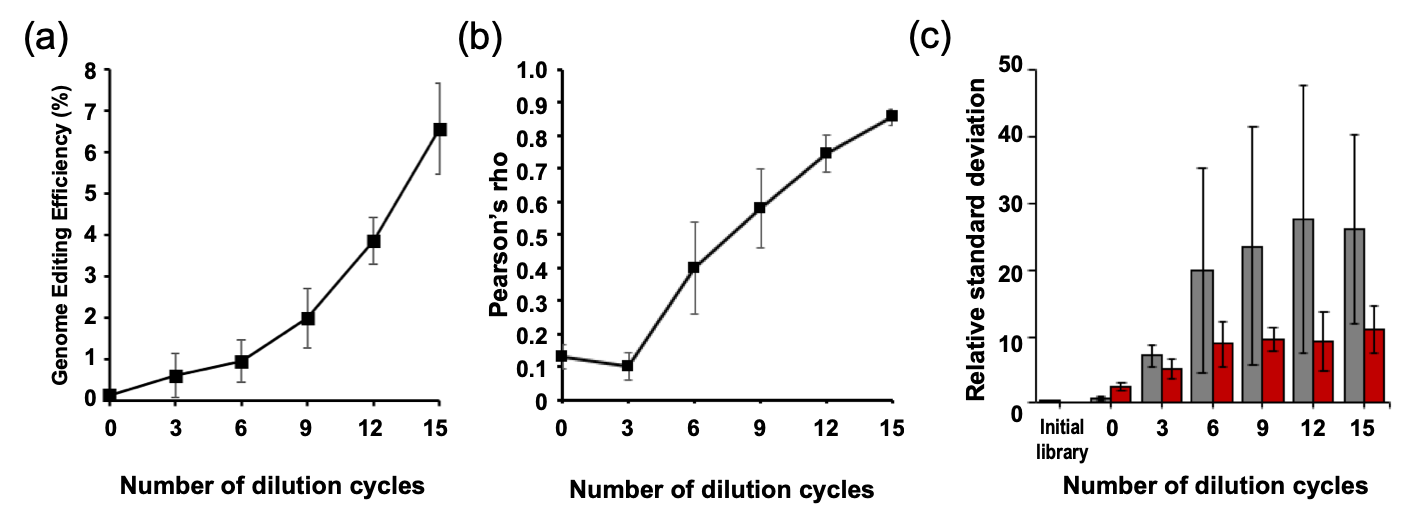

**Figure S3** | Results of the degenerate sequence experiment. (a) Genome editing efficiency, (b) correlation coefficient, and (c) bias of each degenerate sequence with increasing number of dilution cycles. Bias was determined as relative standard deviation. Each graph was plotted based on two independent experiments, and all error bars indicate standard deviation.

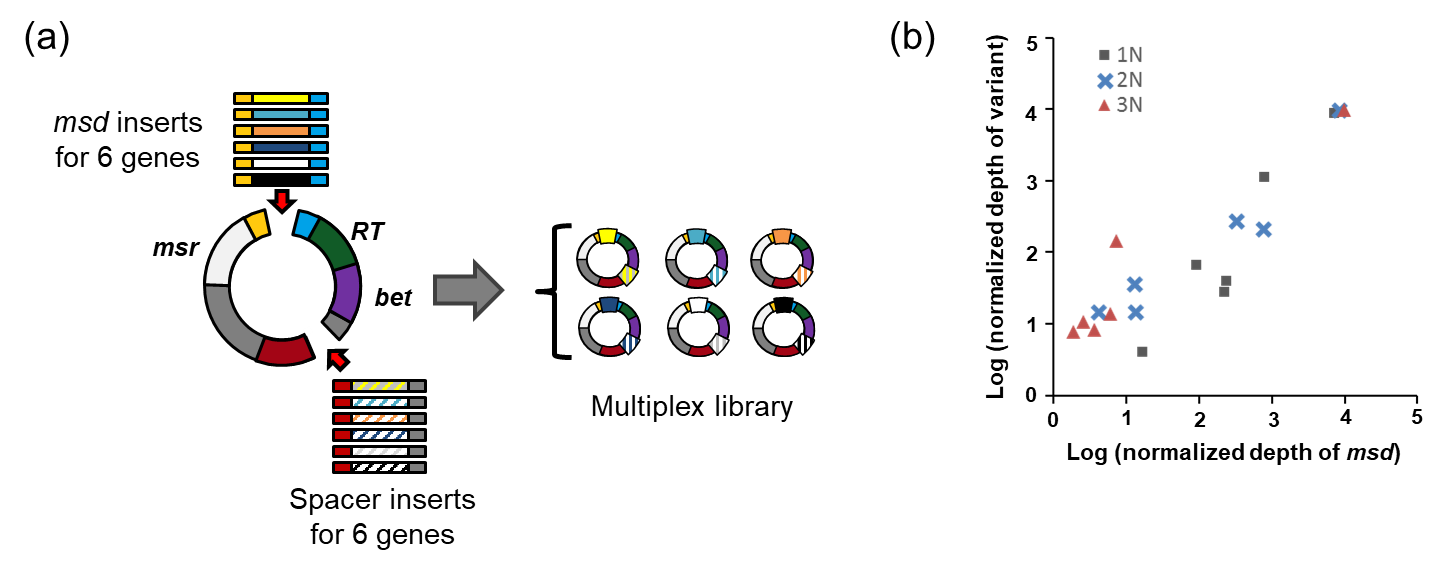

**Figure S4** | Multiplex engineering experiment protocol and results. (a) Schematic illustration of the procedure for multiplex library generation targeting six genes in the *E. coli* genome. The *msd* and matched spacer inserts were cloned individually and subsequently mixed to construct the multiplex library. (b) Correlation of *msd* and variant depth (Pearson correlation coefficient [ρ]=0.96, n=3). Squares, crosses, and triangles denote n=1, n=2, and n=3, respectively.

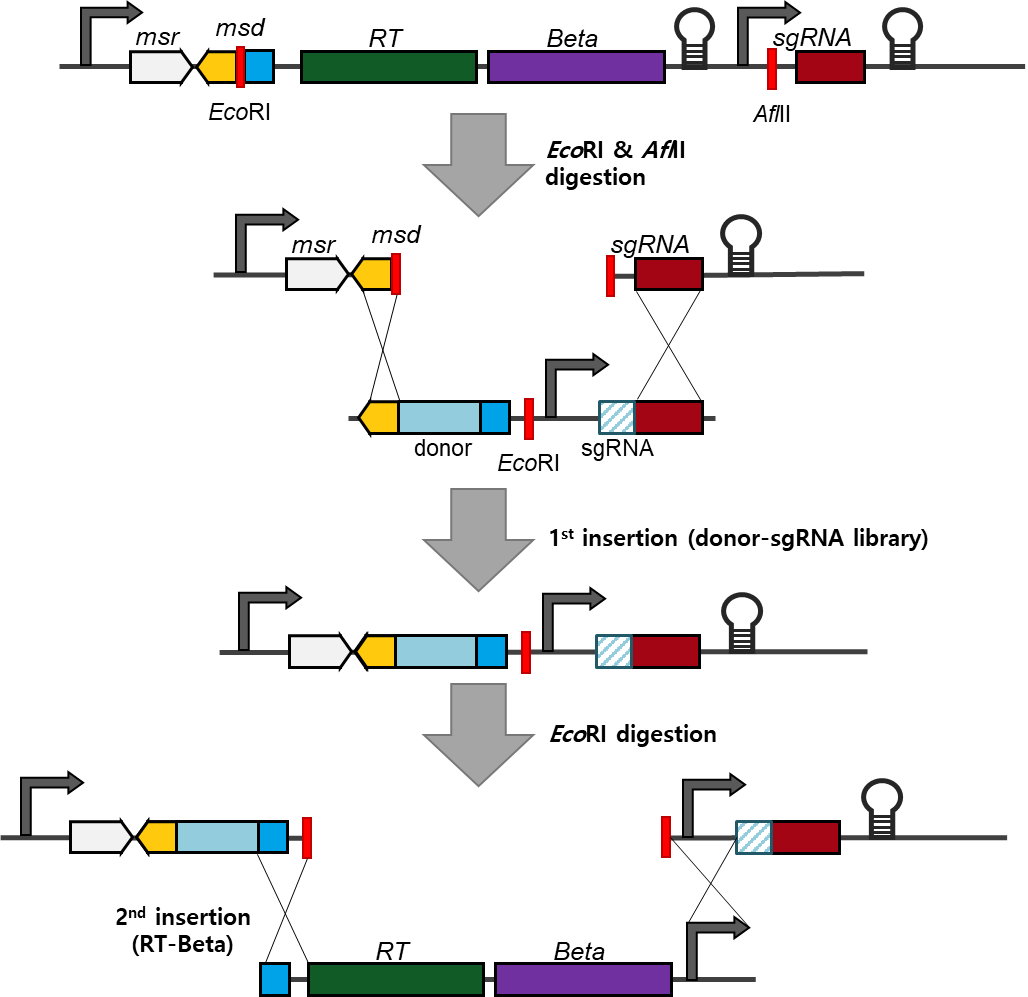

**Figure S5** | Schematic illustration of the strategy used for construction of the genome-scale library. The plasmid library was constructed by following two steps: (1) first insertion: library insertion into *Eco*RI- and *Afl*II- digested pRC_blank_02; (2) second insertion: insertion of RT-beta fragment into the *Eco*RI-digested plasmid produced in the first insertion.

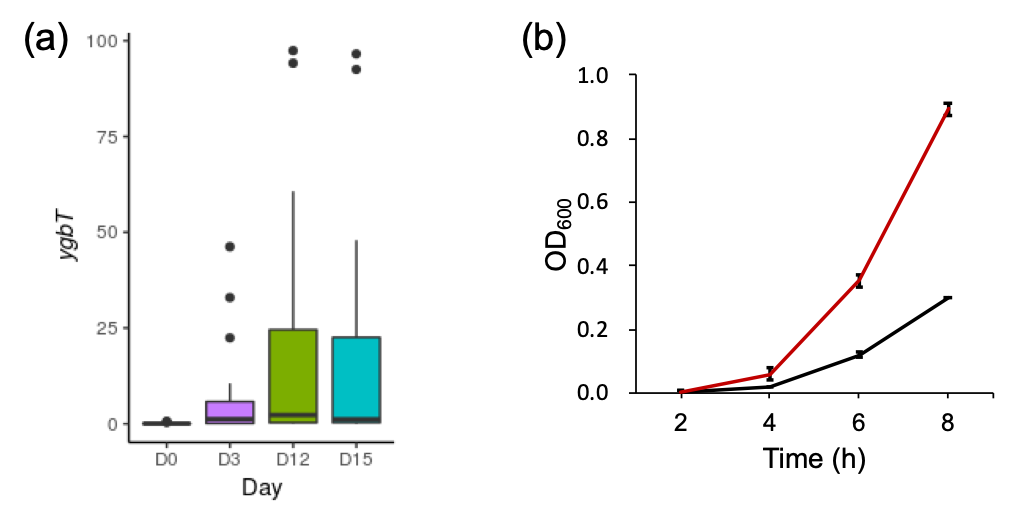

**Figure S6** | Results of the genome-scale experiment. (a) Abundance of *ygbT* at Day 0, 3, 12, and 15. The center line corresponds to the median; the edges of the box correspond to the first and third quartiles; the whiskers represent 1.5× the interquartile range; and the black dots indicate outliers. (b) Growth curves of cells bearing *ygbT*-targeting plasmid (red) or control plasmid (pRC_blank_02) (black). Each graph was plotted based on three independent experiments, and error bars indicate standard error of the mean.

**Supplementary Tables**

**Supplementary Table 1** | List of the parts and sequences used in the study.

**Supplementary Table 2** | List of plasmids used in the study.

**Supplementary Table 3** | List of strains used in the study.

**Supplementary Table 4** | Clonal validation of engineered subpopulation in the N6 oligo-based experiment. Genomic loci and the *msd* region in singly picked colonies were validated by Sanger sequencing. All engineered sequences in the genomic loci were consistent with their *msd* region.

**Supplementary Table 5** | Target information.

**Supplementary Table 6** | List of oligos used in the study.

**Supplementary Table 7** | Calculated fold-change in the chloramphenicol-treatment experiment. One-way ANOVA and Tukey post-hoc analysis were performed based on this data.

**Supplementary Table 8** | Results of one-way ANOVA.

**Supplementary Table 9** | Results of Tukey post-hoc analysis.

**Supplementary Table 10** | Clonal validation of the engineered subpopulation in the chloramphenicol selection experiment. The *cat* locus was validated in singly picked colonies by Sanger sequencing.

**Supplementary Table 11** | List of oligos used for the genome-scale library.

**Supplementary Table 12** | Results of ANOVA for the ordinary least-square model.
